# VIEW-MOD: A Versatile Illumination Engine With a Modular Optical Design for Fluorescence Microscopy

**DOI:** 10.1101/660332

**Authors:** Bei Liu, Chad M. Hobson, Frederico M. Pimenta, Evan Nelsen, Joe Hsiao, Timothy O’Brien, Michael R. Falvo, Klaus M. Hahn, Richard Superfine

## Abstract

We developed VIEW-MOD (Versatile Illumination Engine With a Modular Optical Design): a compact, multi-modality microscope, which accommodates multiple illumination schemes including variable angle total internal reflection, point scanning and vertical/horizontal light sheet. This system allows combining and flexibly switching between different illuminations and imaging modes by employing three electrically tunable lenses and two fast-steering mirrors. This versatile optics design provides control of 6 degrees of freedom of the illumination source (3 translation, 2 tilt, and beam shape) plus the axial position of the imaging plane. We also developed standalone software with an easy-to-use GUI to calibrate and control the microscope. We demonstrate the applications of this system and software in biosensor imaging, optogenetics and fast 3D volume imaging. This system is ready to fit into complex imaging circumstances requiring precise control of illumination and detection paths, and has a broad scope of usability for a myriad of biological applications.

## 1. Introduction

Fluorescence microscopy (FM) is one of most powerful techniques in biological and biomedical research. A handful of FM techniques have been developed to accelerate acquisition speed, improve resolution, reduce background and/or reduce phototoxicity. The excitation strategies of FM can be divided into three major categories: widefield, point-scanning, and light-sheet illumination. In a widefield configuration collimated light exits the objective illuminating the entire sample. We are specifically interested in total internal reflection microscopy (TIRF) which effectively channels the illumination light to within a few hundred nanometers of the substrate. With point-scanning, the objective focuses a collimated light source onto a diffraction limited point, which is raster scanned across the sample to render an image. In light-sheet illumination a thin sheet of excitation light is created either by focusing one axis of the illumination source or rapidly scanning a focused beam along a single direction. Scanning the light sheet across the sample can then create a 3D volume. In the following, we briefly introduce these three modes to highlight the advantages of a combined system.

Total internal reflection fluorescence microscopy (TIRF) is a widefield illumination technique that suppresses out-of-focus fluorescence by adopting evanescent waves for excitation. TIRF has been broadly used for single molecule detection and for studying dynamic events near the membrane. Data acquisition, however, suffers from the interference fringes due to the coherent nature of lasers. Several groups have reduced the coherence of the laser light prior to illumination either through a multi-mode fiber [1] or spinning/vibrating a diffusor [2–3]. Other groups have performed fast scanning of the azimuth angle during a single exposure through a rotating wedge [4], a steering mirror [5–7], an Acoustic Optical Deflector [8], or a Digital Micromirror Device [9]. The former may lead to loss of light polarization that is critical in many applications. Uniform TIRF illumination can alternatively be achieved by forming a ring of laser illumination at the back-focal-plane (BFP) of the objective. The 360-degree in-plane (from all sides), polarized illumination will generate a radially symmetric evanescent field exhibiting a flattop intensity profile across the field-of-view [10]. This technique is commonly referred to as variable angle or spinning TIRF (vaTIRF or spinTIRF). Commercially available vaTIRF systems such as iLas2 from Roper Scientific and TILL/ FEI’s iMIC with the Polytrope illuminator are expensive and lack the flexibility to combine other imaging modalities.

Point illumination is also a powerful technique as it allows researchers to selectively illuminate a portion of the sample without perturbing other areas. This technique is particularly useful in the field of optogenetics to precisely activate or inhibit particular proteins with precise spatiotemporal resolution. However most of the experiments were carried out with either a separate photoactivation module [11] or a commercialized confocal microscope [12]. Implementing both point and widefield in the same light path is non-trivial, yet would allow for further techniques to be employed in separate paths of an optical system.

Light sheet fluorescence microscopy (LSFM) is an increasingly popular technique for imaging biological specimens at high spatiotemporal resolution with reduced background and phototoxicity; therefore LSFM is well suited for long-term 3D imaging of both single cells and tissue [13]. Numerous variations of light sheets (Gaussian, Bessel beam, line Bessel sheet, Airy beam, etc.) have been demonstrated in the past, each providing a unique means of illuminating a thin slice of a given sample [13]. The most notable of these techniques is lattice light sheet, where a two-dimensional optical lattice is used to create thin, structured illumination sheets that are dithered during a single exposure [14]. Traditionally LSFM requires two objective lenses, one for illumination and one for detection, placed at a 90-degree angle relative to each other such that the light sheet from the illumination source coincides with the imaging plane of the detection objective [14–21]. Such systems can be less user-friendly than conventional microscopes. High numerical aperture (N.A.) immersion objectives are almost precluded from LSFM applications due to the geometry configurations and the physical dimensions of the objectives. Both our research group and others have implemented LSFM in a single-objective system [22–25]; here we present how we can combine single-objective LSFM into a microscope design that allows for other FM techniques.

Each of the above imaging modalities has distinct advantages; a system capable of each technique is then ideal for studying various biological processes. Other groups have previously developed multi-modality microscopes that combine two separate super resolution methods [26], multiple light sheets [27], or epifluorescence and TIRF [28–30]. These systems are however not adaptable to all the illumination techniques we have outlined. Here, we present the first microscope design to our knowledge capable of widefield (particularly vaTIRF), point scanning, and LSFM. Our design is compact, modular for customization, cost effective, and built upon a conventional inverted microscope, allowing it to be implemented in most laboratories. The different illumination modes can run independently or be switched in a single experiment, allowing one to combine or change the imaging modality to fit the experiment at hand. The microscope is operated with an open-source software package (available upon request) to calibrate and control the system, which is independent of the microscope automation software.

## 2. System Design

Here we present a versatile optics design that provides precise control of the translation, tilt, and shape of illumination light at the back focal plane (BFP) of the objective and/or the sample plane, as well as axial control of the imaging plane through a series of electrically tunable lenses (ETL) and scanning mirrors (SM) (Fig. 1). Our design is conceptually divided into a series of modules (Fig. 1, dashed box 1-5), each with a specific purpose for shaping and controlling the light through the system. Modules 1-4 govern the illumination pathway while module 5 lies in the detection pathway. Through using some or all of the modules available, our system can implement a variety of imaging modalities. Here, we outline the theory and purpose of each module, with the reader directed to Figure 1 for the optical layout.

**Fig. 1.**
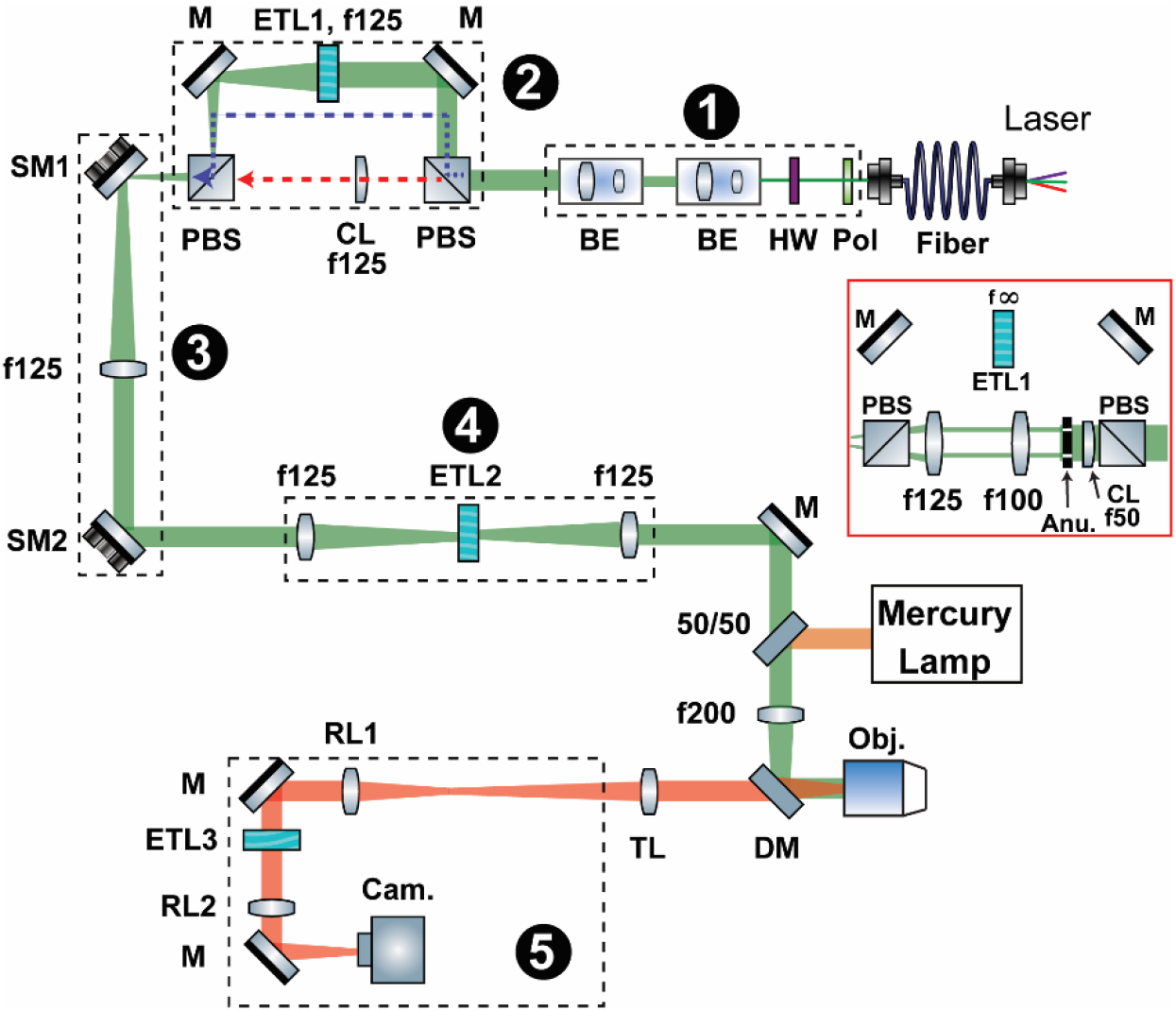
Schematics of the system. Linearly polarized lasers are expanded, collimated and exit Module 1. The light propagates to Module 2 through either pathway 1 (blue dashed-line) for TIRF and point-scanning or pathway 2 (red dashed-line) for LSFM, depending on the orientation of its polarization axis. For TIRF, ETL1 is adjusted to 125 mm effective focal length to focus the beam onto SM1, resulting in a collimated light beam coming out the objective. For point-scanning ETL1 is set to ~ 0 volts, equivalent to f=∞ (flat plate). For LSFM, we either put a 125 mm cylindrical lens to create a Gaussian light sheet or use a combination of cylindrical lens and annulus to create a Line Bessel Sheet (LBS). Steering Mirrors 1 and 2 (SM1 and SM2) are optically conjugated with the specimen plane and the Back Focal Plane (BFP) of the objective, respectively. ETL2 provides axial scanning of the illumination light. In the detection module (Module 5), the image is projected to the camera through a relay lens group (RL1 and RL2) with ETL3 placed in the center (mimicking Module 4) to achieve an adjustable image plane. Schematic was generated with GW Optics component library (http://www.gwoptics.org/ComponentLibrary/).

### Module 1: Beam expansion and polarization modulation

Module 1 controls the size and polarization of the illumination light. Multiple linearly polarized lasers are coupled into a single-mode, polarization-maintaining fiber (PM-S405-XP, Thorlabs). The light from the fiber is expanded and collimated through commercially available collimators (GBE05-A and BE052-A, Thorlabs). This properly sizes the incident beam for a given application, such as filling the back aperture of the objective lens for point illumination to obtain the tightest focus. The polarization axis of the light exiting the optical fiber is then adjusted by either rotating a half-wave plate (AHWP05M-600, Thorlabs) or through a liquid crystal retarder (LCC1111T-A, Thorlabs). A polarized beam splitter (PBS) redirects the light either to light path 1 (LP1) or LP2 of module 2 to switch between a circular beam cross section or a cylindrical (light sheet) cross section. The PBS determines the orientation of the polarization of the beam, and the tunable waveplate governs the amplitude of the light going into each arm of the following module. We note that the system is designed such that when using light sheet illumination, the polarization vector of the illumination light is aligned with the plane of the light sheet. In some cases, it may be useful to switch the polarization state of the illumination beam at the specimen. That can be accomplished by adding a second waveplate after module 2. The use of a tunable filter allows computer control of these operations.

### Module 2: Beam shaping

The beam-shaping module accommodates two light paths for different applications. The switching between LP1 and LP2 is achieved by altering the polarization direction of the incident laser using module 1. This can be computer controlled by employing the liquid crystal retarder. LP1 retains the incident circular cross section, while allowing for quick transitions from widefield illumination to point illumination with ETL1 (EL-16-40-TC, Optotune). When the ETL is set such that the focal length approaches infinity, light is not deflected resulting in a collimated beam at the BFP of the objective; the objective focuses the light to a diffraction-limited point at the sample. This will be useful, for example, for photoactivation employed below. The focal length of the ETL can be adjusted such that the beam is focused at the BFP of the objective, resulting in a wide, collimated illumination at the sample. This will be employed for our TIRF imaging. The response and settling times of the ETL are 5 ms and 25 ms respectively, allowing for dynamic switching of imaging modalities on the order of exposure times for biological imaging. LP2 is used to shape the illumination light to be cylindrical in cross section, which is frequently used in light sheet imaging. LP2 contains an optic, typically a cylindrical lens, to converge one of the transverse axes of the light while keeping the second axis collimated, creating a cylindrical pattern. These two paths are then rejoined through a second polarized beam splitter. Having two illumination geometries in a single system allows one to employ drastically different illumination techniques (e.g. LSFM and point scanning) on a single sample.

### Module 3: Beam steering

Module 3 controls the steering of the illumination light both at the BFP of the objective and at the sample plane using two fast steering mirrors (SM1,2) (OIM 101 1”, Optics In Motion LLC). SM1 is optically conjugate to the BFP of the objective while SM2 is optically conjugate to the sample plane. The former controls the position of the beam at the sample plane and the latter adjusts the tilt at the sample plane. SM1 and SM2 are driven by voice coils and are each able to control two axes of tilt in a single mirror. This allows a more compact design for optical conjugation with a single mirror compared with using galvo-scanners, which typically needs two mirrors in a 4f configuration. While galvo-scanners are notably faster, SM1 and SM2 can execute a 1 mrad step in under 5 ms, providing enough speed for sub-second volumetric imaging.

### Module 4: Axial beam scanning

Module 4 provides control of the axial position of the illumination light without physical displacement of the objective lens. Keeping the objective motionless is ideal as it reduced small motions of the sample that are coupled to objective movement. We use a 4f system to relay the beam and position a second ETL (ETL2) (EL-16-40-TC, Optotune) midway between the second and third relay lens (conjugate to the BFP of the objective). Adjusting the focal length of ETL2 subsequently scans the focus of the illumination light axially [31]. This can be especially useful for both LSFM and point scanning to ensure that the tightest focus of the illumination light is at the desired axial location in the sample.

### Module 5: Axial image plane scanning

Module 5 is an identical 4f system to that of module 4, lying in the detection path. Similarly, we place an ETL (ETL3) (EL-16-40-TC, Optotune) midway between two relay lenses. Adjusting the focal length of ETL3 then shifts the axial position of the imaging plane [24,32,33]. One can dynamically manipulate the focus of an image during an experiment without moving the objective lens or the illumination light. It should be noted that large scale changes of the optical power of ETL3 will (de)magnify the image [17,34]. Other groups have used an ETL for axial control of the imaging plane by placing the ETL directly behind the detection objective [17,34–37], however using ETL3 in a 4f system makes it easier to access and adjust as well as minimizes the (de)magnification effects [32,33]. We ensure that ETL3 lies flat (its optical axis is normal to the table) to avoid gravitational effects which would distort the Optotune lens.

Next, we will demonstrate how these proposed modules can implement vaTIRF, point scanning, and light-sheet illumination.

## 3. System Implementation

### 3.1 vaTIRF and point scanning

To implement vaTIRF and point scanning, we use modules 1-3. Module 1 sizes the illumination light and adjusts the polarization such that module 2 directs the light along LP1, establishing a circular cross section for the beam. Switching between widefield and point illumination is achieved by changing the current applied on ETL1, while beam steering is controlled as described in module 3. Tilting SM2 (conjugate with the sample plane) translates the beam at the BFP of the objective, resulting in a change in tilt at the sample plane, see Fig. 7(a) in Appendix A. SM2 can then be used to scan the beam to form a focused circle at the BFP of the objective. Similarly, tilting SM1 (conjugate with the objective BFP) will cause a tilt of the beam at the objective BFP and therefore a lateral movement of the beam at the sample plane, see Fig. 7(b) in Appendix A. SM1 and SM2 are controlled by four channels of analog voltage supplied by a National Instrument (NI) DAQ board (PCIe-6323). The SMs need to be calibrated to establish the correspondence between the control voltages and the beam position at either the BFP of the objective or the sample plane. To calibrate SM2, a small CCD camera (UI-3370CP Rev. 2, IDS) was placed at the location of the BFP of the objective. The voltages for both axes of SM2 were scanned from −10 V to 10 V in increments of 1 V. An image was taken at each voltage pair to record the beam position (Fig. 8(a) in Appendix B), and a 2D surface function (‘poly55’ in MATLAB) was fitted to the beam positions and the corresponding voltage pair (Fig. 8(b), 8(c) in Appendix B). To calibrate SM1, we used an autofluorescent slide (92001, Chroma) on the microscope specimen stage. SM1 voltages were scanned from −10 V to 10 V in increments of 1 V with an image acquired at each voltage pair (Fig. 8(d) in Appendix B) by the imaging sCMOS camera (ORCA-Flash4.0 V3, Hamamatsu). Another 2D surface was fitted to establish the mapping between the control voltages of SM1 and the beam position at the sample plane (Fig. 8(e), 8(f) in Appendix B).

For widefield imaging, spinning the beam during exposure minimizes laser self-interference artifacts. To achieve TIRF the scanning radius should be large enough so that the incident angle is greater than the critical angle. The relationship between scanning radius and incident angle can be illustrated by geometrical optics, see Fig. 8(g) in Appendix B. The penetration depth (pd) can be calculated as:

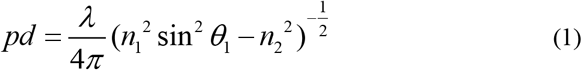

where pd is determined only by the wavelength of the laser (λ), the refractive index of the substrate (n1) and that of the media (n2), and the incident angle *θ*_1_ [38]. To determine the incident angle for each wavelength we placed a ruler next to the microscope a distance, D, away from the objective. When the laser beam is deviated r away from the center of the objective the incident angle is denoted as *θ*_1_. From the Fresnel equation, we can calculate the incident angle as

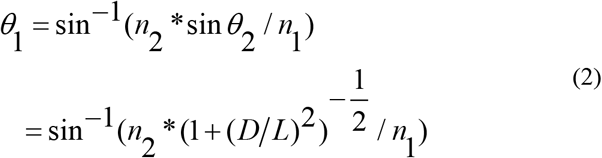

where *L* and *θ*_2_ are as defined in Fig. 8(g) in Appendix B.

We plotted the scanning radius, r, against the incident angle and fitted a linear function to determine the scanning radius at which we reach the critical angle (Fig. 8(h) in Appendix B). While the scanning radius remains the same for different wavelengths of incident light, the penetration depth varies according to the above equation (Fig. 8(i) in Appendix B). For multicolor TIRF imaging, it is essential to ensure that all lasers are collimated through the objective lens; we can compensate for chromatic aberration from different wavelengths by adjusting ETL1. Equivalently, if we choose to maintain the same penetration depth for each illumination color, we can use a lookup table to program the scanning radius for each color.

The microscope, camera, filter wheels, auto-focus, and lasers are controlled with either Metamorph or Micromanager. A home-built MATLAB program is used to control the NI DAQ board and ETL1 and to run the calibration procedure. The synchronization between camera exposure and laser beam steering is achieved by connecting the TTL out signal of the imaging camera with one of the Programmable Function Interface (PFI) lines on the NI DAQ board, as shown in Fig. 9 in Appendix C.

### 3.2 Light-sheet illumination

To implement LSFM, we employ modules 1-6. Module 1 properly sizes the beam and rotates the polarization state such that the light passes through LP2, the cylindrical cross section path of module 2, and the polarization vector is in the plane of the light sheet. We use a Line Bessel Sheet (LBS) because it provides an extended depth of field for a given waist size when compared to a Gaussian light sheet. To create an LBS we use a cylindrical lens (LJ1567RM-A, Thorlabs) to focus one axis of a collimated light source at the location of the annulus (R1DF200, Thorlabs), resulting in two coherent bands of light that propagate through the remaining optical path. The annulus is optically conjugate to the BFP of the objective. When the two coherent bands pass through the objective lens (UplanSAPO 60x/1.2 W, Olympus), they interfere with each other resulting in a Line Bessel Sheet [19]. The measured lateral and axial FWHM of the LBS is 771 nm (Fig. 2(a)) and 14.8 μm (Fig. 2(b)) respectively, both of which match well with the theoretical value [39,40]. One could also easily create a Gaussian light sheet by placing a 100 mm cylindrical lens (LJ1567RM-A, Thorlabs) before SM1 in place of the current LBS pathway.

**Fig. 2.**
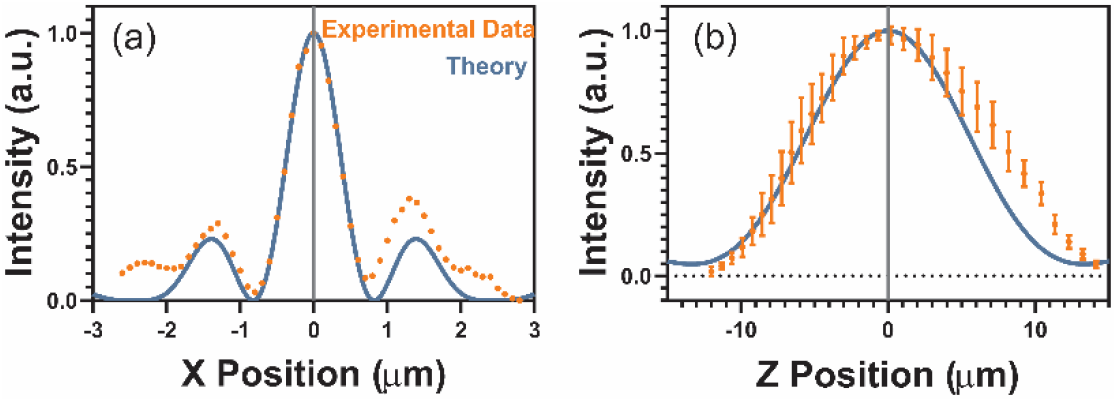
Profile of LBS. (a) Lateral profile of the LBS. A 100 nm fluorescent bead was stepped laterally in 100 nm increments by a piezo stage; intensity of the bead was measured in FIJI for each position. (b) Axial profile of LBS. 100 nm fluorescent beads were imaged as the LBS was stepped axially by ETL2 in 2 mA increments. Intensity of 7 beads was tracked across the scan in FIJI, error bars are standard deviations in intensity. Orange represents measured intensity from the fluorescent beads; blue represents a theoretical profile [39,40].

Modules 3 and 4 provide complete control of all three spatial positions of the illumination light while module 5 allows us to adjust the axial position of the imaging plane. We use a home-built LabView program that controls two NI DAQ boards (PCIe-6323 and PCI-6723); one controls voltages to the SM1 controller and ETL3 controller (TR-CL180, Gardasoft) while the other controls an acousto-optic tunable filter (AOTF:AOTnC-400.650-TN, Opto-Electronic) in our light engine to gate the laser light. We use the Optotune software to control the current applied to ETL2 which subsequently varies the optical power.

As noted earlier, SM1 is conjugate to the BFP of the objective. Tilting SM1 translates the LBS in x-y at the sample plane. For volumetric imaging, we need to control only the translation of the LBS in the x direction (perpendicular to the long axis of the light sheet). We can then use a linear combination of the two voltage controls to SM1 to create a single voltage control parameter that translates the LBS in the x direction. To calibrate SM1, we used a fluorescent slide to visualize the LBS. We incrementally stepped the SM1 control voltage while taking an image at each step; the center of the LBS was determined for each image and plotted against its corresponding voltage. A line was fit to the resulting plot to determine a conversion of SM1 voltage to LBS translation, see Fig. 10 (a) in Appendix D. Scanning SM1 allows us to translate the LBS laterally and collect volume images, and the calibration provides a means of calculating the voxel size. To calibrate ETL2, we again used a fluorescent slide to visualize the LBS. The objective lens was incrementally stepped and ETL3 was used to readjust image plane back to the coverslip. ETL2 was then used to lower the LBS back to the fluorescent substrate. This provided a calibration for how ETL2’s current adjusts the axial position of the LBS, see Fig. 10 (b) in Appendix D. Adjusting ETL2 ensures that the thinnest portion of the LBS is always located in our sample without manually adjusting the objective height. The Gardasoft controller for ETL3 is programmed to the linear analog mode (0 V – 10 V) and outputs a current based on a closed-loop temperature feedback system. To calibrate ETL3, we place a grid (R1L3S3, Thorlabs) on the microscope and manually step the objective lens position. ETL3 is used to bring the grid back into focus, providing a calibration of ETL3 voltage with the imaging plane position, see Fig. 10 (c) in Appendix D. Imaging plane location, light sheet position and AOTF are synchronized through an NI DAQ board.

## 4. Results

### 4.1 vaTIRF allows homogeneous FRET imaging with reduced photobleaching

Epifluorescence FRET imaging of biosensors in living cells benefits from ratio imaging, in which the FRET signal is divided by the donor signal to normalize for the effects of varying path length through the cell, or variations in biosensor distribution. TIRF imaging largely eliminates the path length effects, but the laser self-interference illumination pattern forces significant corrections to obtain images suitable for ratio imaging [41]. A previous study showed that vaTIRF could produce homogeneous FRET images [42]. We tested here whether vaTIRF could enhance the accuracy of biosensor ratio images by examining the migration of Mouse Embryonic Fibroblasts (MEF) stably expressing a single chain FRET biosensor that reports RhoA activation [43]. Cells were sequentially imaged with epifluorescence, fixed-angle TIRF, and vaTIRF (Fig. 3). Ratio images indicated regions of higher RhoA activation during cell migration (Visualization S1, S2), showing activation at the leading and retracting edges of the cell, as in previous studies [43].

**Fig. 3.**
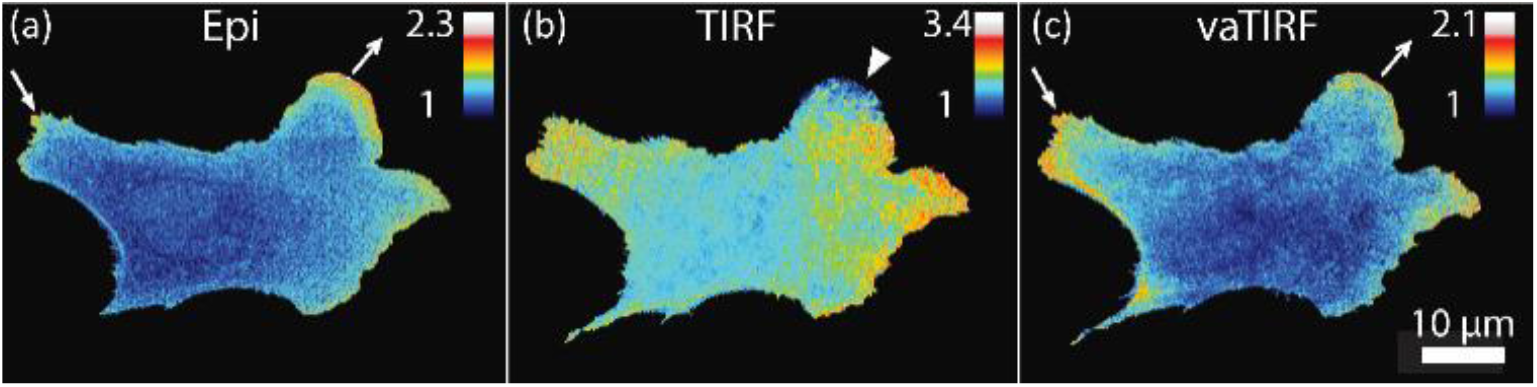
vaTIRF imaging of RhoA activity using a FRET biosensor. Mouse Embryonic Fibroblast (MEF) cell stably expressing a RhoA biosensor [43] imaged sequentially with epifluorescence illumination (a), fixed angle TIRF (b) and vaTIRF (c) to compare different imaging modes. FRET/Donor ratio images from epifluorescence and vaTIRF showed RhoA activity in cell protrusions (outward pointing arrow) and retractions (inward pointing arrow), while uneven illumination with fixed angle TIRF produced spurious activation maps (e.g. inward pointing arrow head).

Fixed-angle TIRF failed to report GTPase activity accurately because it was difficult to correct for the sinusoidal illumination pattern produced by interference, especially when the signal was faint or reduced to noise in some regions. The biosensor’s dynamic range was similar in epifluorescence and vaTIRF, accurately reflecting the low and high activation states of the biosensor, see Fig. 11 (a) in Appendix E. Photobleaching of the biosensor was greatly reduced when using vaTIRF rather than traditional epifluorescence, see Fig. 11 (b) in Appendix E, enabling imaging over longer periods or more images for greater kinetic resolution.

### 4.2 Localized photoactivation of Rac1 promoted cell protrusion and formation of adhesions

We can transition from vaTIRF to point-scanning by changing the current sent to ELT1. To highlight the importance of this capability in optogenetics, we combined PA-Rac1, an optogenetic tool that induces membrane protrusion upon blue light stimulation [11], with the adhesion marker Paxillin. We used vaTIRF to image a MEF migrating on a fibronectin-coated coverslip, while using point-illumination to locally photoactivate PA-Rac1 (Fig. 4, Visualization S3). We observed mostly static adhesions with small constitutive membrane movements before photoactivation (< 5 min), whereas during localized photoactivation of PA-Rac1 (5 – 15 min) we observed increased membrane protrusion near the irradiation site, followed by the formation of new adhesions. After this photoactivation period (15 – 20 min) the cell membrane retracted and the newly formed adhesions dissipated.

**Fig. 4.**
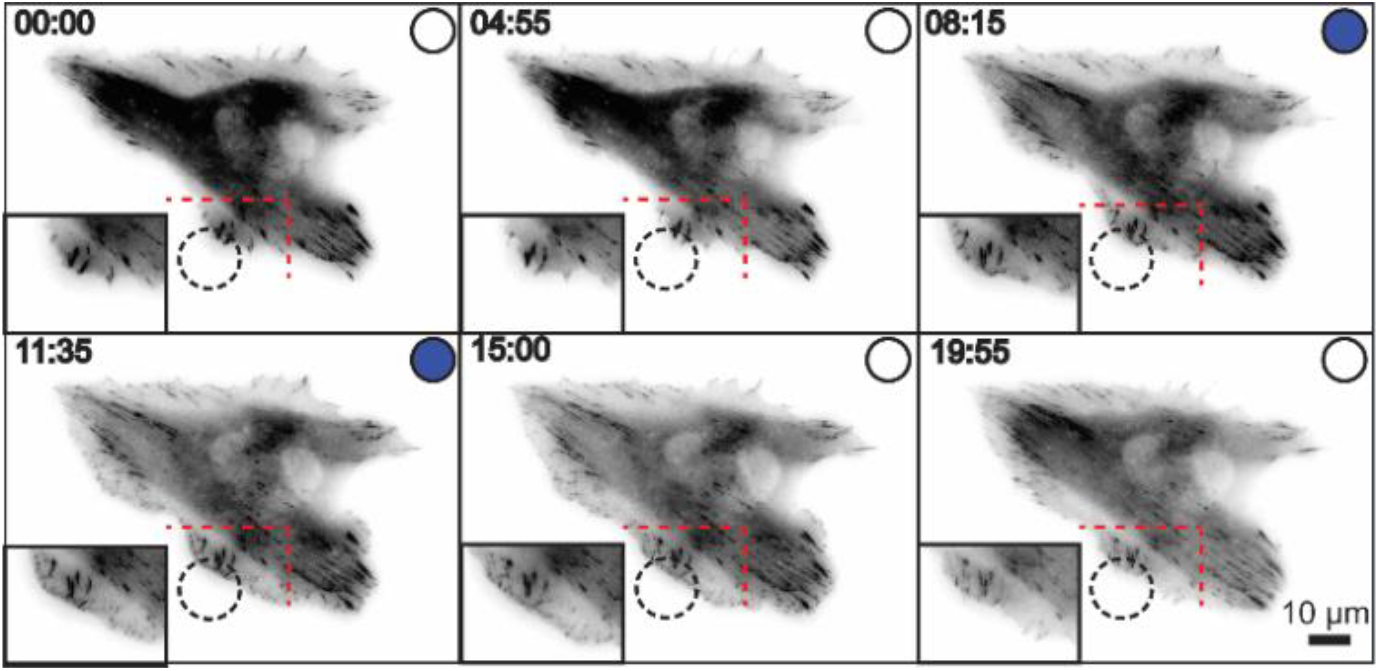
Combined vaTIRF imaging of Paxillin and localized photoactivation of PA-Rac1. A Mouse Embryonic Fibroblast (MEF) cell stably expressing mVenus-photoactivatable Rac1 (PA-Rac1) [11] was transiently transfected with mCherry-Paxillin, an adhesion marker. Paxillin was imaged using vaTIRF for a total of 20 min and the cell was locally irradiated (dashed circle) with pulsed blue light for 1 second every 5 seconds (500 μW) between the 5 and 15 min mark (blue-filled circle on the top of each panel indicates photoactivation). Localized photoactivation at the cell edge led to protrusion with the formation of small adhesions, followed by cell retraction and dissipation of adhesions.

### 4.3 Horizontal and vertical LSFM of highlights organization of heterochromatin and euchromatin

We employed two LSFM methods, Horizontal Light Sheet (HLS) and Vertical Light Sheet (VLS), to image chromatin of cell nuclei (Fig. 5). For both methods we introduced a small (180 μm) right-angle reflective prism (8531-607-1, Precision Optics) attached to a 6 degree-of-freedom mount, and lowered the mirror adjacent to the cell of interest [25]. For HLS imaging, the LBS reflected off the mirror and illuminated a single x-y plane. SM1, ETL2, and ETL3 were used to appropriately position the LBS and the imaging plane. For VLS imaging, the LBS propagated vertically out of the objective and through the cell, illuminating a single y-z slice. The objective was raised until the imaging plane intercepted the reflective optic. This rotated the imaging plane from x-y space to y-z space. As the objective was continually raised, the imaging plane stepped through the cell in X until it was aligned with the illuminated slice. As a demonstration of these techniques, we used both HLS and VLS to sequentially image orthogonal planes of a COS-7 cell nuclei stably expressing HaloTag-H2B labeled with Janelia Fluor 549 (JF549) (Fig. 5 (c), (d)).

**Fig. 5.**
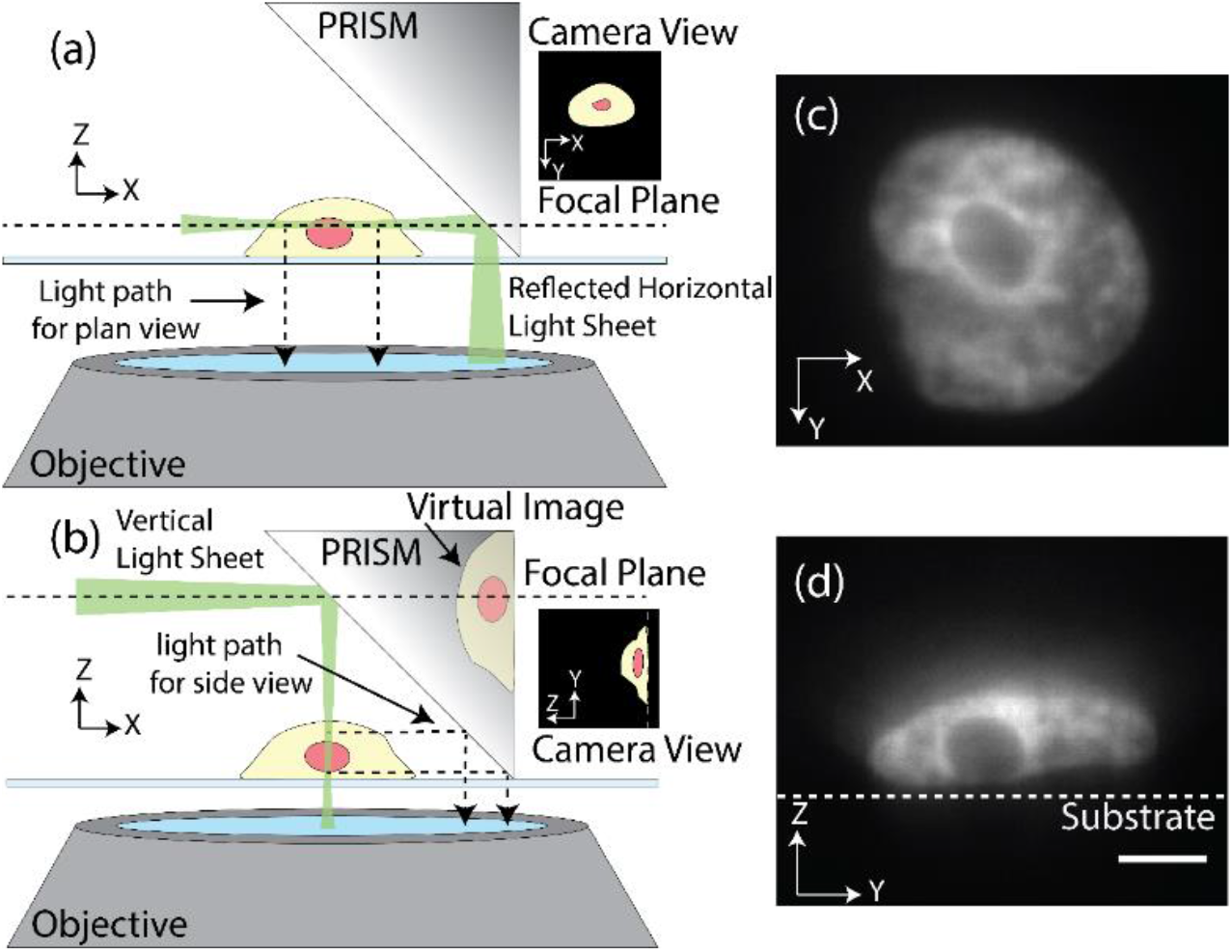
LSFM of orthogonal planes of heterochromatin and euchromatin. (a) Imaging setup for Horizontal Light Sheet. The light sheet emerges from the objective and is reflected by a right-angle prism. The image plane is matched (by module 5) to the plane of the reflected light sheet. (b) Imaging setup for Vertical Light Sheet. The light sheet emerges from the objective and creates a vertical slice through sample. The right-angle prism creates a virtual side-view image of the vertical slice. Horizontal (c) and vertical (d) light sheet images of the same COS-7 nucleus expressing HaloTag-H2B labeled with Janelia Fluor 549. Scale bar = 5 μm.

Transitioning from HLS to VLS is straightforward, fast, and can be done entirely through software. It is important to note that resolution is not affected when the imaging plane is rotated by the mirror [25]. Because of LSFM’s ability to illuminate a single plane, we obtained images with low background and high signal-to-noise of orthogonal planes of the same nucleus. We observed that the light collection efficiency for a given illumination intensity and exposure time using HLS is approximately double that of using VLS. For highly sensitive imaging techniques such as FRET it is beneficial to use HLS imaging due to the larger signal-to-noise. However, AFM studies are better served with VLS imaging as it allows the user to take fast, high resolution images of the plane in which the AFM tip is applying force without the need for taking whole stacks [44–46]. Because of the lack of background fluorescence, we could discern heterochromatin (high H2B density) from euchromatin (low H2B density) within the nucleus. Specifically, we observed that a thin ring of heterochromatin surrounds the nucleolus. Visualizing chromatin structure is important in understanding chromosome territories and gene expression [47].

### 4.4 Live-cell LSFM volumetric imaging of filopodia dynamics

This microscope design with LBS illumination is also capable of fast volumetric imaging by incrementally stepping the light sheet across a sample and continuously matching the imaging plane to the position of the light sheet. In a single-objective system, it is slightly more complicated due to the coupling of the light sheet position and imaging plane. Even though both HLS and VLS can be used to take volume images, here we chose VLS because it requires one less degree of freedom to control through software than HLS.

We used SM1 to step the LBS in increments laterally in the x-axis through the cell, and ETL3 was adjusted such that at each step the image was in focus. Every time the mirror was brought down next to a cell of interest, the spacing of the cell and mirror was different and the angle of the mirror relative to the substrate was slightly varied due to the mechanical micrometer control mechanism. For each cell we performed an initialization scan (Fig. 12 in Appendix F) yielding a lookup table of the ETL3 voltage value that provided the best focus for each LBS position in the volume scan. With this lookup table we performed fast volume scans of RAW 264.7 macrophage cells stably expressing HaloTag-F-Tractin labeled with JF549 (Fig. 6, Visualization S4). Each slice was taken in 10 ms (5 ms exposure, 5 ms transition), and each volume consisted of 75 slices meaning that a single volume image was taken in 0.75 s; this is comparable to recent two-objective systems [14,17,19]. Because of the speed of our system, we were able to observe both formation and movement of filopodia on the timescale of several seconds. These timescale dynamics are fundamental to processes such as phagocytosis and cell migration [48,49], and hence our system is well suited to study mechanisms in which macrophages migrate and engulf foreign particles.

**Fig. 6.**
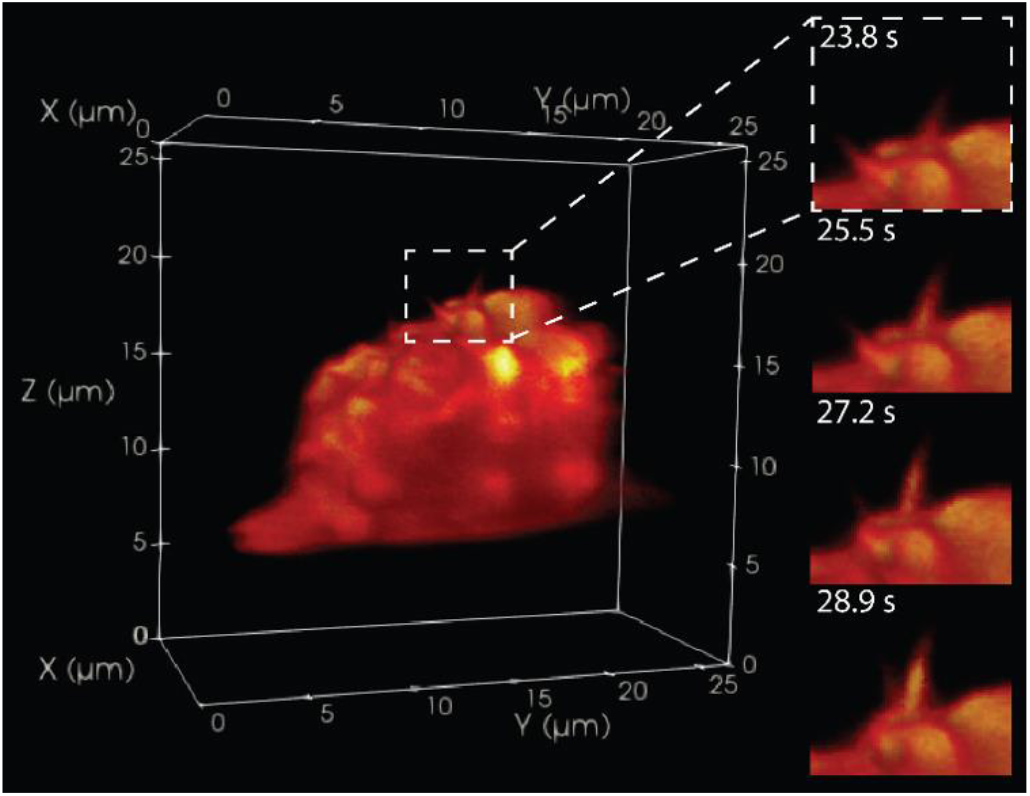
Vertical light sheet volume imaging showed filopodia formation and dynamics. Selected frame from a volumetric movie (Visualization S4) of a RAW 264.7 macrophage cell expressing HaloTag-F-tractin labeled with JF549. Each volume consists of 75 slices each taken in 10 ms (5 ms exposure, 5 ms transition time) for a 0.75 s total volume acquisition time with 100 ms delay between each volume. Total volume size is 12.5 μm x 26.7 μm x 25.9 μm, voxel size is 167 nm x 106 nm x 106 nm. We observed formation and movement of individual filopodium on second time scales.

## 5. Discussion

We have demonstrated the first microscope design capable of vaTIRF, point-scanning, and LSFM, the first localized photoactivation combined with vaTIRF, and the first single-objecive LSFM imaging of orthogonal planes of a single cell nucleus. Further novelty of this instrument, however, lies in its broad scope of applicability. We have outlined a table of various advanced microscopy techniques our system can implement as well as the required modules to do so (Table 1 in Appendix G). Many of the available techniques involve scanning of different illumination patterns. By scanning the incident angle in vaTIRF, the penetration depth of the evanescent field is modified. Additionally stepping the imaging plane in sequence with the incident angle, one can axially section the range of the evanescent field and construct 3D renderings of near-surface interactions [50]. One could also adjust the incident angle such that it is slightly below that of TIRF to achieve HILO microscopy [51]. Point illumination can be further extended by introducing two-photon microscopy, which allows for femtoliter focal volume precision [52]. Provided use of the proper laser and sample, our design can produce precise two-photon point illumination in 3D which allows for photoablation, photoactivation, and fluorescence recovery after photobleaching (FRAP) studies.

Our system has the ability to both laterally and axially scan a light sheet, which when combined with the rolling shutter functionality of an sCMOS camera can produce x-y or y-z images with minimal background [16,53]. We also have the capability to create light sheets by dithering a focused beam at a higher frequency than the imaging rate. Our modules allow for precise shaping of the focused beam, meaning that a scanned beam could be tailored to image sub - cellular structures or multi-cell systems depending on the interest of the research group. Scanned beams could be used in our system for both x-y and y-z imaging as well as volumetric imaging similar to what we previously demonstrated [24]. A focused beam could also be incrementally stepped through a sample perpendicular to the image plane to create a spatially structured illumination pattern for HiLo microscopy [54]. Finally, our system is capable of super-resolution imaging through either stochastic optical reconstruction microscopy (STORM) or photo activated localization microscopy (PALM) in conjugation with TIRF or LSFM. Both STORM and PALM rely upon on-off switching fluorophores and an illumination source with low background [55]. Provided the proper sample and software development (much of which is openly available), our microscope’s TIRF and LSFM can be used for these super-resolution techniques.

## 6. Conclusions

It is essential for a microscopy system to both visualize and control the distribution and activity of target proteins. Optical systems are often designed around a single microscopy technique and even though such systems have drastically advanced our ability to investigate single cell dynamics, the focus on individual techniques can limit the scope of available research questions. Here we have described the design for a single, compact versatile optics system capable of vaTIRF, point-scanning and LSFM imaging. This design is cost-effective and built upon a conventional inverted microscope, making it easily implementable in any laboratory with the latter. Furthermore, the open-source software can be expanded to include other techniques due to the complete control of illumination light and imaging plane location. In its current implementation, we showed that vaTIRF provides superior quality for ratiometric imaging compared to conventional TIRF, with significantly reduced photobleaching than epifluorescence. We further combined this technique with localized activation of an optogenetic tool, opening new venues of research where one is able to locally induce activation of signaling enzymes and observe their downstream effects. Finally, we used LSFM to image x-y and y-z cross sections of chromatin as well as capture fast volumetric images of filopodia dynamics. This microscope design can be easily adapted to serve a user’s needs, make use of the ever-growing number of light-sensitive tools being developed, and is minimally perturbing to normal cell physiology thus allowing great potential in opening new venues of research and answering important biological questions.

## Supporting information

Visualization J1

Visualization J2

Visualization J3

Visualization J4

## Appendix A: Ray tracing while scanning SM2 and SM1 for vaTIRF and point-scanning

**Fig. 7.**
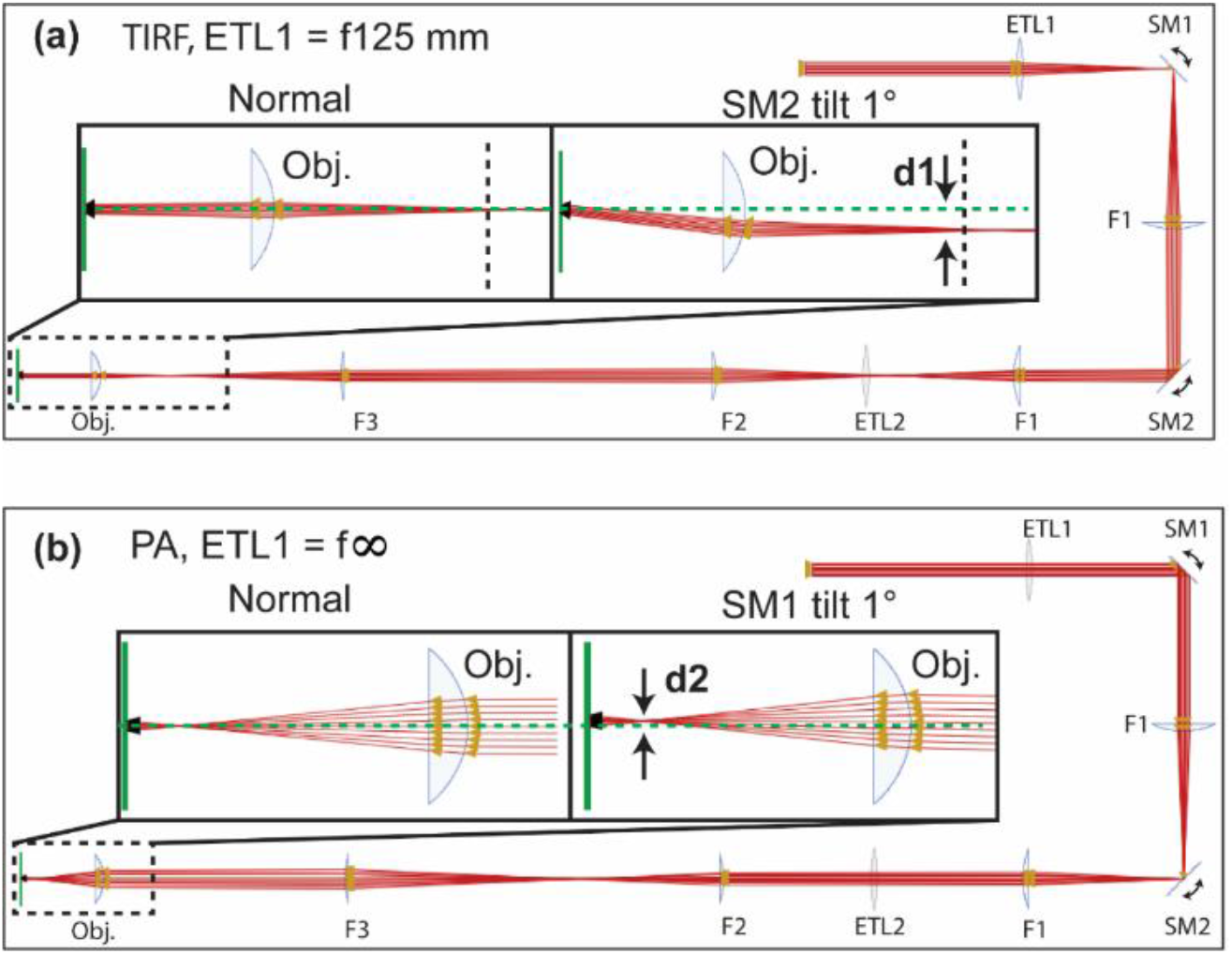
Ray tracing is done with *OpticalRayTracer* (*https://arachnoid.com/OpticalRayTracer/*). (a) For TIRF, ETL1 is set to a value with 125 mm effective focal length. The laser is focused on the BFP of the objective, which is indicated by the arrows in the insert. Tilting of SM2 will cause beam displacement at the BFP of the objective ***d1***, but due to conjugation, the beam will not shift on the sample plane. **(b)** For PA, ETL1 is set to 0. The beam is collimated before entering the objective and focused on the sample plane, which is indicated by the arrows in the insert. Due to conjugation, tilting of SM1 will not shift the beam at the BFP of the objective but change the incident angle, which will move the focused point on the sample plane ***d2***

## Appendix B: Calibration of vaTIRF and point-scanning

**Fig. 8.**
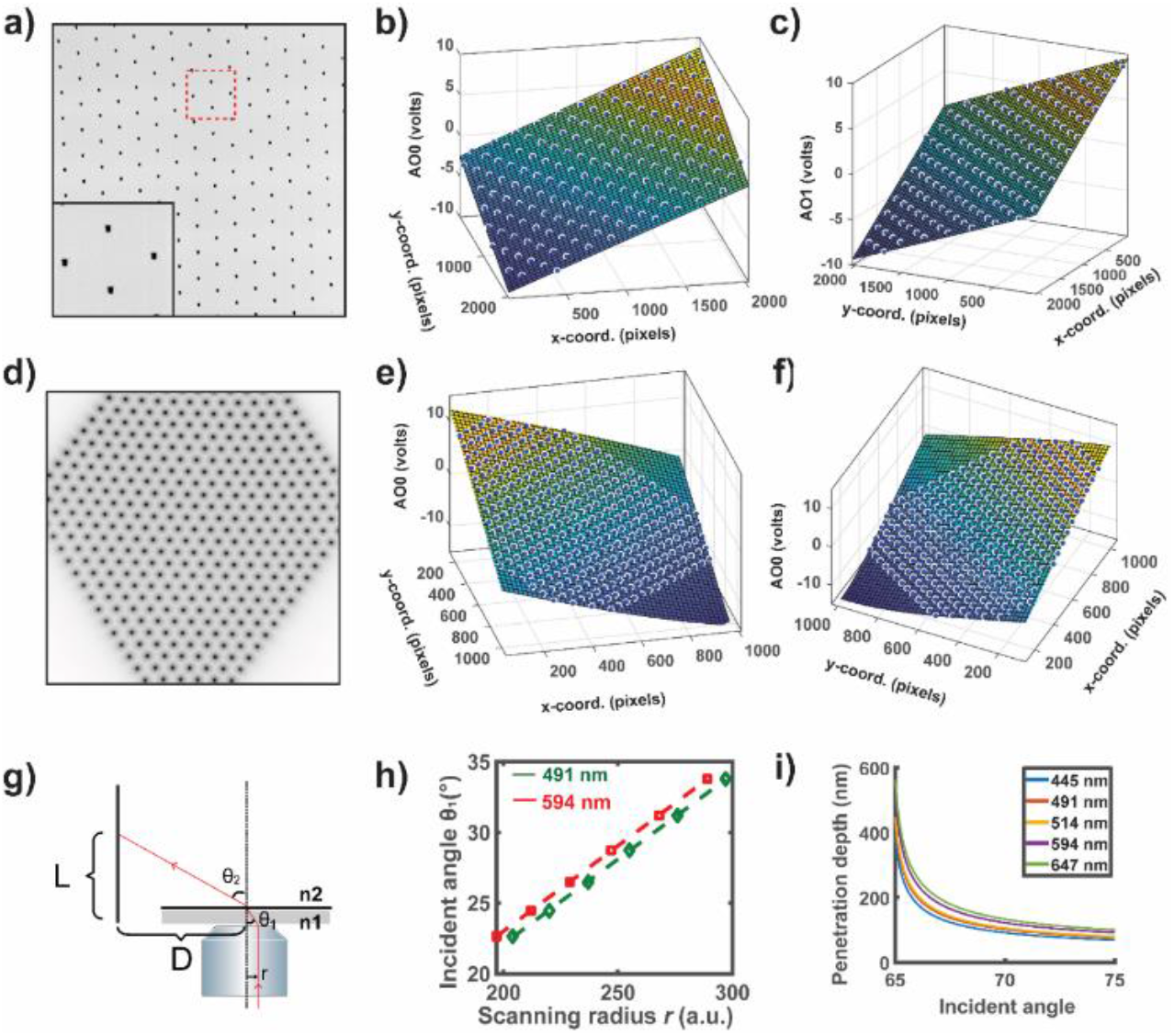
(a) Maximum projection of 441 images taken with the camera mounted at the BFP of the objective for SM2 calibration. The insert indicated the overlay of all found beams. (b, c) Fitted surface to map beam positions (x, y)_BFP_ at the BFP of the objective to voltage pairs (vx, vy)_BEP_. (d) Maximum projection of 441 images taken with the sCMOS cameras used for SM1 calibration. (e, f) Fitted surface to map beam positions (x, y)_sample_ at the sample plane to voltage pairs (vx, vy)_sample_. (g) Illustration of linking scanning radius, *r*, with penetration depth. *Θ_1_* is the incident angle. (h) The relationship between scanning radius and incident angle (or penetration depth) is wavelength dependent. To establish the correspondence, the laser beam (for instance 491 nm and 594 nm lasers here) was scanned with a pre-defined radius, *r*. The beam position on the ruler, *L*, was recorded and converted to the incident angle, and then plotted against the scanning radius and fitted with a line. (i) Plot of the scanning radius against penetration depth for different wavelength.

## Appendix C: Software design and hardware triggering for vaTIRF and point-scanning

**Fig. 9:**
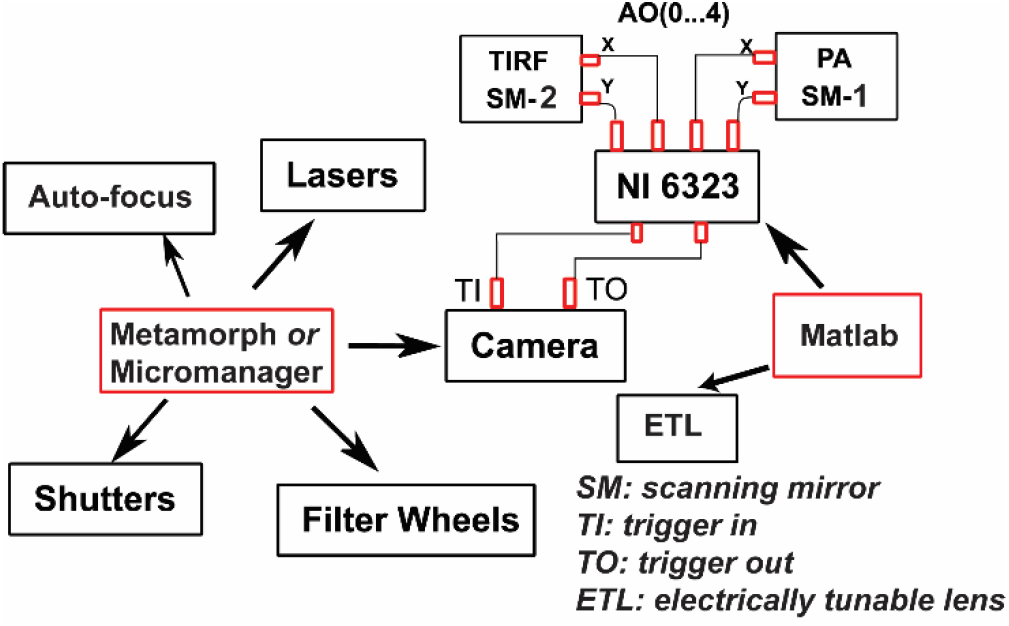
We decouple mirror scanning and ETL control from general microscope automation so the software can be easily combined with any given system. We use Metamorph and/or μManager to control other microscope components such as cameras, lasers, shutters, filter wheels and others. Home-built software was used to control two scanning mirrors through NI Data Acquisition Boards and ETLs. The synchronization between camera exposure and mirror scanning was achieved through the trigger in and trigger out function of cameras.

## Appendix D: Calibration of SM1, ETL2, and ETL3 for LSFM

**Fig. 10.**
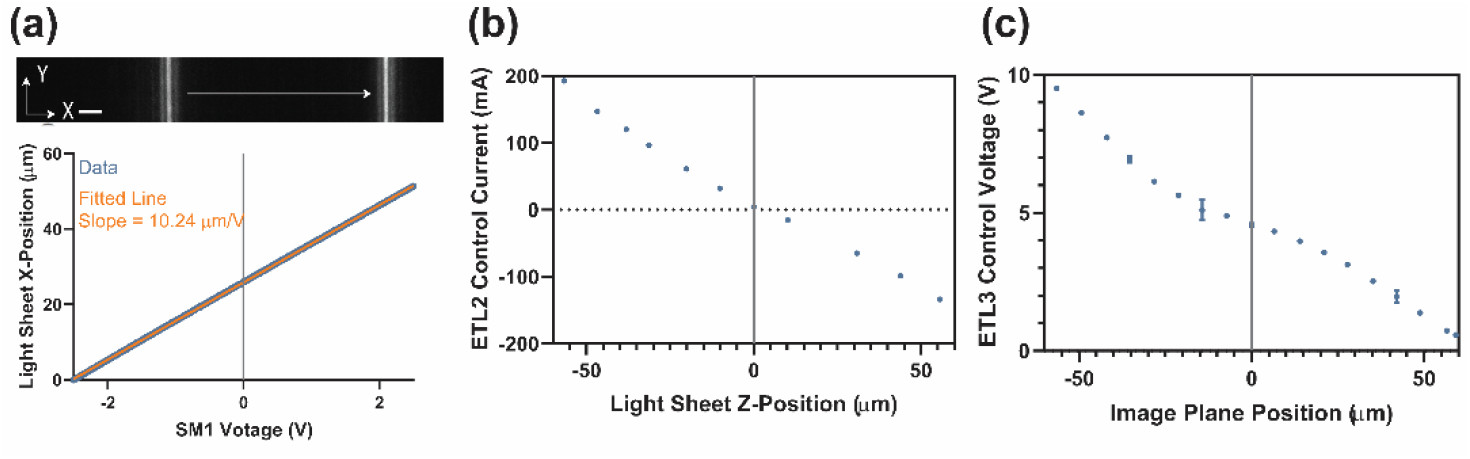
(a) Calibration of SM1 showing the light sheet x-position versus voltage applied to SM1 as well as a linear fit to this data. The slope of the linear fit provides us a conversion from SM1 voltage to scanning distance, allowing us to understand and properly set our voxel size during volume imaging. Scale bar = 5 μm. (b) Calibration of ETL2 showing the current applied to ETL2 versus the light sheet z-position. (c) Calibration of ETL3 showing the voltage applied to ETL3 versus the axial position of the image plane. Error bars for (b) and (c) are 95% confidence intervals, but are often smaller than the visualization of the data points.

## Appendix E: Dynamic Range and photobleaching comparison of vaTIRF and Epifluorescence

**Fig. 11:**
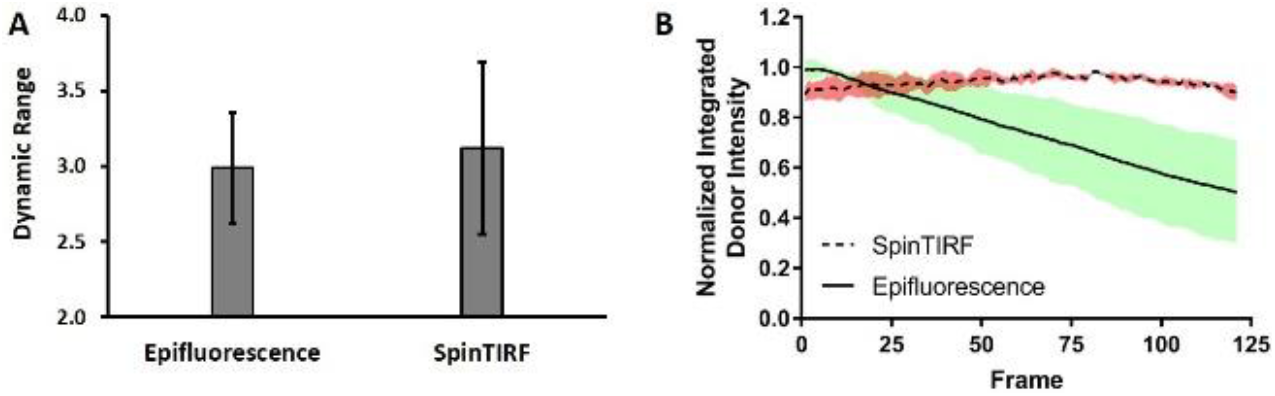
(A) Dynamic range comparison of a RhoA single-chain biosensor using data from Epifluorescence (n=12) and SpinTIRF (n=12) imaging, which showed no significant difference. (B) Corrected photobleaching curve of the RhoA biosensor stably expressed in MEF cells, using either widefield (n =3) or vaTIRF (n =4) (200 μW for Epifluorescence, 500 μW laser power at the stage, 15 minutes time course, with same number of images and exposure time). Photobleaching curves were corrected for the power difference between the two imaging modes. There was a significant reduction in photobleaching with vaTIRF illumination.

## Appendix F: Initialization scan procedure for volumetric imaging

**Fig. 12.**
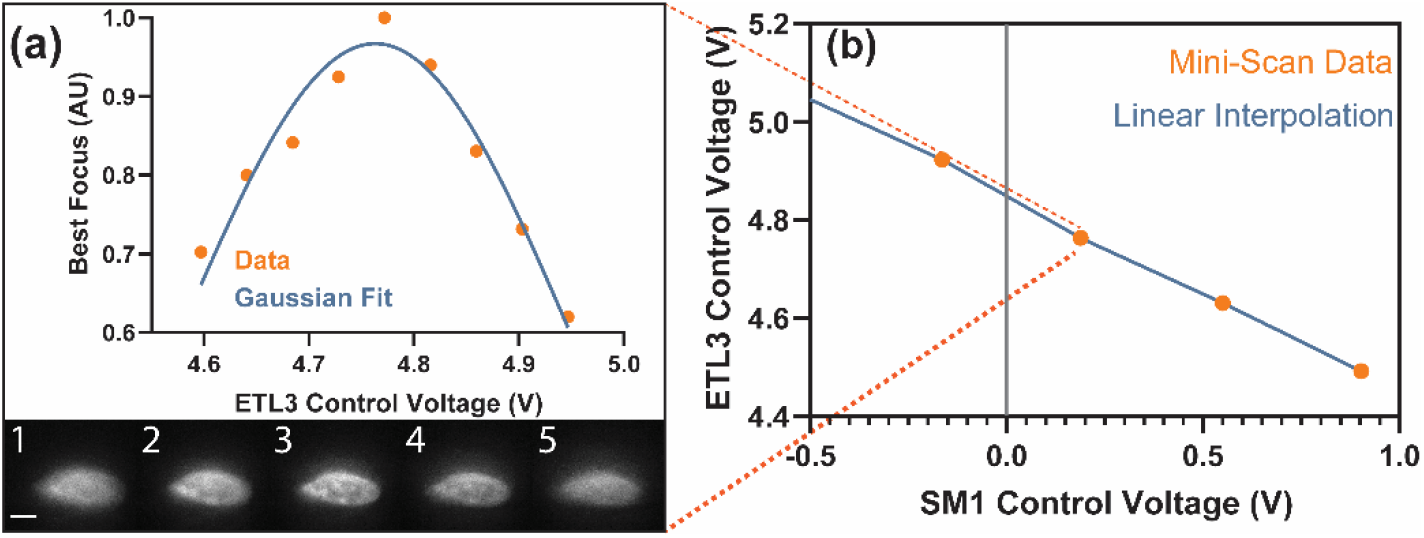
(a) A sample mini scan of a COS-7 nucleus expressing HaloTag-H2B labeled with Janelia Fluor 549 used in the initialization scan where the LBS is fixed to a specific location and the ETL3 voltage is varied about a suggested voltage of best focus. The Tenegrad method [15,56]is used on each image from the mini scan to provide a metric for image quality. A Gaussian is fit to the mini-scan data to provide the optimal ETL3 voltage for a given light sheet position. (b) A sample lookup table that provides the best ETL3 voltage for a given light sheet position. Each orange data point is generated by a mini scan procedure. A linear interpolation is done between data points and is discretized based upon the number of slices per volume.

## Appendix G: Expanded list of imaging modalities for VIEW-MOD

**Table 1.** We provide an extensive, but not necessarily complete, list of imaging modalities in the scope of VIEW-MOD’s capabilities. The modules required for each modality and are listed, only further software development would be required for implementation. *Module 1: Beam expansion and polarization modulation. Module 2: Beam shaping. Module 3: Beam steering. Module 4: Axial beam scanning. Module 5: Axial image plane scanning.

| Imaging Modality | Description | Required Modules* | Reference |
| --- | --- | --- | --- |
| Single-Objective Light Sheet Fluorescence Microscopy (2D & 3D) | As described in main text | 1-4 (2D), 5 (3D) | Present Work |
| Localized Point Illumination (2D & 3D) | As described in main text. This could be extended to control the z-position of the beam with use of module 5. | 1-3, 4 (3D) | Present Work |
| TIRF (variable angle or traditional) | As described in main text | 1-3 | Present Work |
| Widefield Epifluorescence | As described in main text | 1-3 | Present Work |
| Volumetric Imaging with vaTIRF | Varying the incident angle subsequently varies the penetration depth of the evanescent field. Scanning the incident angle in connection with the imaging plane allows for z-sectioning within the full range of the evanescent field. Image stacks can be taken and reconstructed to form volumetric images. | 1-3,5 | [50] |
| Laterally swept LSFM with rolling shutter | A light sheet propagates vertically from the objective lens and scans laterally either through the sample (y-z light sheet plane, x-y imaging plane) or across the prism (x-y light sheet plane, y-z imaging plane). The rolling shutter of a sCMOS camera is matched to the position of the light sheet as it is scanned to create high SNR images. | 1-5 | [53] |
| Single-Objective Scanned Beam LSFM (2D & 3D) | A focused beam is scanned at a higher frequency than the image acquisition rate to create a light sheet. This light sheet can be used similarly to as we have described in the main text. | 1-4 (2D), 5 (3D) | [24] |
| Axially Swept LSFM with rolling shutter | A light sheet propagates vertically from the objective lens and scans laterally either through the sample (y-z light sheet and imaging plane) or across the prism (x-y light sheet and imaging plane). The rolling shutter of a sCMOS camera is matched to the position of the waist of the light sheet as it is scanned to create high SNR images. | 1-4 | [16] |
| Fluorescence Recovery After Photobleaching (FRAP) | A focused light beam of high intensity bleaches a localized section of a sample, and the sample is then imaged and the recovery of the fluorescent intensity at the location of the bleaching is monitored. This lets a user understand the timescale of diffusion of the fluorescent molecules in the sample. | 1-4 | [57] |
| 2-Photon 3D Localized Excitation | With an appropriate laser and sample choice, our system can implement 2-photon microscopy wherein one could achieve a focal volume on the femtoliter order for the illumination light. This could be used to perform precisely localized photoactivation or laser ablation in 3D. | 1-3, 4 (3D) | [52] |
| HILO Microscopy | A beam is focused and translated laterally at the back focal plane of the objective such that the incident angle at the sample is just below that of TIRF, allowing a highly inclined beam to pass through the sample for improve signal-to-background over widefield. | 1-3 | [51] |
| STORM/PALM | STORM and PALM are super-resolution techniques that rely upon on-off switching of fluorophores. With the proper sample, our microscope design is capable of these techniques using either TIRF or LSFM. | 1-4 (TIRF), 5 (LSFM) | [55] |

## Appendix H: Cell culture

### H1. vaTIRF and Point Scanning

Stable Mouse Embryonic Fibroblast (MEF) cell lines for the single-chain RhoA FRET biosensor and PA-Rac1 were generated and cultured as previously described [43]. Three days before imaging, cells were washed with fresh medium to remove doxycycline. For both biosensor and photoactivation experiments cells were seeded in fibronectin-coated coverslips 2-4 hours prior to imaging, and maintained in regular culture media. PA-Rac1 stable cell lines were kept in the dark after doxycycline removal. Cells were imaged in F-12 Ham’s media (Caisson Labs) supplemented with 5% FBS (Gemini) and 1 mM HEPES (Gibco). MEFs stably expressing mVenus-PA-Rac1 were transiently transfected with pEGFP-C1 mCherry-Paxillin (Clontech) using TransIT-X2 (Mirus) 24 hours prior to imaging. Cell were washed with fresh media ~4 hours after transfection.

### H2. LSFM

Murine macrophage cells (RAW 264.7) were transfected with HaloTag-F-tractin gene using Fugene HD (Promega), and grown in phenol free DMEM F12 with 10% FBS. Similarly, COS-7 cells (a gift from the Liu Lab at Janelia Research Campus) stably expressing HaloTag-H2B were grown in the same medium. For experiments, both cell types were plated sparsely on 55 kPa polyacrylamide (PA) gel pads that had been coated with fibronectin.

Gel pads were prepared on 40 mm circular #0 coverslips (Fisher Scientific) to minimize reflections from a glass surface onto the side view mirror. Coverglasses were UV cleaned and treated with APTES (Amino propyl triethoxy silane; 1% in toluene as vapor). 55 KPa gels were prepared from standard protocols, with the addition of 1% polyacrylic acid before polymerization. Once polymerization was initiated in a small volume of acrylamide with 10% APS, 10 μL was placed at the center of two coverglasses, and a 22 x 22 mm coverslip placed rapidly on top of each. These squares were treated with hexamethyldisilazane vapor after UV cleaning to develop a hydrophobic, non-adherent surface. A small weight was placed on the coverslip to promote spreading of the acrylamide in the few seconds before it gels.

After 5 minutes of polymerization, DI water was placed around the upper coverslip and a scalpel blade was used to gently pry up the upper coverslip. The gels were then allowed to dry slightly so that 1 cm cloning cylinders could be affixed to the gel with Dow High Vacuum grease. Then, 150 μL of fibronectin (human; 10 μg/ml in PBS) was placed within the cloning rings for 5 minutes before removal. The gels were allowed to dry for 10 minutes in a biosafety hood. The gels were then exposed to the bio-hood UV light for 5 minutes to sterilize them, and 200 μL of either PBS or medium was added until the cells were ready.

After cells had spread on the gels, and 30 minutes before an experiment, a 2-5 μL of a 10^-5^ M solution of Janelia Fluor 549 (JF549) halo ligand (a gift from the Lavis Lab at Janelia Research Campus) in PBS was added to 200 μL of media in the cloning rings. The JF549 solution was then replaced twice with warmed media before imaging. Imaging was carried out in the same medium the cells were grown in.

## Appendix I: Image Acquisition and Processing

### I1. Biosensor Imaging and Processing

MEFs expressing the RhoA biosensor were imaged using a 445 nm laser line or a Mercury Arc lamp (ET430/24X, Chroma with a 10% Neutral Density Filter) with Donor (FF01-475/42, Semrock) and FRET (ET535/30M, Chroma) images acquired for 500 ms (Dichroic mirror zt440/514/561/640rpc-UF2). Movies and images were processed using a MatLab package previously developed for FRET biosensor imaging (available at http://www.hahnlab.com/tools/software.html). Briefly, Donor and FRET movies were corrected for uneven illumination (shade correction), background subtracted, thresholded for exclusion of all non-cell signal, intensity corrected for photobleaching and finally FRET/Donor was calculated. Power from both the laser line and the Mercury arc lamp was measured by placing the power meter head (Thorlabs S121B) at the microscope revolver. For the direct comparison between Epifluorescence, fixed-angle TIRF and vaTIRF the laser power was matched to the lamp power required to acquire FRET/Donor movies in Epifluorescence (500 μW); the same applies for dynamic range comparisons. Photobleaching curves were obtained by integrating the total Donor intensity (after shade correction, background subtraction and thresholding) for every frame, corrected for power difference (200 μW for vaTIRF, 500 μW for Epifluorescence), and finally divided by the area of the cell for vaTIRF but not for Epifluorescence; Changes in cell shape over the duration of the movie correlate linearly with the signal in vaTIRF but not in Epifluorescence.

### I2. Combined photoactivation and vaTIRF

mCherry-Paxillin images were acquired every 5 sec using a 561 nm laser line (Coherent OBIS 514nm LX 40 mW; emission filter FF01-600/37, Semrock) and PA-Rac1 was photoactivated with a 445 nm laser line between the 5 and 15 min mark for 500 ms at every frame (Dichroic mirror zt440/514/561/640rpc-UF2).

### I3. LSFM

COS-7 cells stably expressing HaloTag-H2B were labeled with JF549 and imaged with a 561 nm laser line (Coherent OBIS 561nm LS 150 mW). Images were acquired with a 200 ms exposure time for both HLS and VLS modes. Laser power at the sample plane was measured with a power meter (ThorLabs PM100D) to be 88 μW.

RAW 264.7 cells stably expressing HaloTag-F-tractin were labeled with JF549 and imaged with a 561 nm laser line. Images were acquired with a 5 ms exposure time and 5 ms transition time, and each volume consisted of 75 images for a total volume acquisition time of 0.75 s. A delay of 100 ms was allowed between each volume image. Laser power at the sample plane was measured to be 27 μW. Volume rendering were produced using ParaView 5.5.2 (Available at https://www.paraview.org/). No image processing was conducted asides from contrasting and mapping intensity to a color and opacity map as to properly visualize the 3D data set.

## Appendix J: Visualizations

### J1. RhoA biosensor imaged using Epifluorescence

Mouse Embryonic Fibroblast (MEF) cell stably expressing a single-chain RhoA biosensor showing RhoA activity during cell motility.

### J2. RhoA biosensor imaged using vaTIRF

Mouse Embryonic Fibroblast (MEF) cell stably expressing a single-chain RhoA biosensor showing RhoA activity during cell motility.

### J3. Combined vaTIRF imaging of Paxillin and localized photoactivation of PA-Rac1

Mouse Embryonic Fibroblast (MEF) cell stably expressing photoactivatable Rac1 (PA-Rac1) [11] transiently transfected with mCherry-Paxillin, an adhesion marker. mCherry-Paxillin was imaged using vaTIRF for a total of 20 min and the cell was locally irradiated (dashed circle) with pulsed blue light for 1 second every 5 seconds (500 μW) between the 5 and 15 min mark (blue-filled circle on the top of each panel indicates photoactivation).

### J4. Volumetric, live-cell imaging of RAW 264.7 cell shows filopodia dynamics

Volumetric movie playing both forwards and then backwards of a live, RAW 264.7 cell expressing HaloTag-F-tractin labeled with JF 549. Various angles are shown with different planes slicing through the volume over the course of the movie to show the internal structure.

## Funding

This material is based upon work supported by the National Institutes of Health (NIBIB Grant Number P41-EB002025), the National Institute of General Medical Science (Grant Number GM-R35GM122596), and the National Science Foundation Graduate Research Fellowship (Grant Number DGE-1650116).

## Acknowledgement

We thank the Liu Lab at Janelia Research Campus for the generous gift of the COS-7 HaloTag-H2B cell line, as well as the Lavis Lab at Janelia Research Campus for the Janelia Fluor 549 Halo Tag fluorescent ligand (JF549).

